# Mutations in a β-group of solute carrier gene are responsible for egg and eye coloration of the *brown egg 4* (*b-4*) mutant in the silkworm, *Bombyx mori*

**DOI:** 10.1101/2021.03.22.436376

**Authors:** Kenta Tomihara, Katsuya Satta, Shohei Matsuzaki, Kazutoshi Yoshitake, Kimiko Yamamoto, Hironobu Uchiyama, Shunsuke Yajima, Ryo Futahashi, Susumu Katsuma, Mizuko Osanai-Futahashi, Takashi Kiuchi

## Abstract

The *brown egg 4* (*b-4*) is a recessive mutant in the silkworm (*Bombyx mori*), whose egg and adult compound eyes exhibit a reddish-brown color instead of normal purple and black, respectively. By double digest restriction-site associated DNA sequencing (ddRAD-seq) analysis, we narrowed down a region linked to the *b-4* phenotype to approximately 1.1 Mb that contains 69 predicted gene models. RNA-seq analysis in a *b-4* strain indicated that one of the candidate genes had a different transcription start site, which generates a short open reading frame. We also found that exon skipping was induced in the same gene due to an insertion of a transposable element in other two *b-4* mutant strains. This gene encoded a putative amino acid transporter that belongs to the β-group of solute carrier (SLC) family and is orthologous to *Drosophila* eye color mutant gene, *mahogany* (*mah*). Accordingly, we named this gene *Bmmah*. We performed CRISPR/Cas9-mediated gene knockout targeting *Bmmah*. Several adult moths in generation 0 (G_0_) had totally or partially reddish-brown compound eyes. We also established three *Bmmah* knockout strains, all of which exhibit reddish-brown eggs and adult compound eyes. Furthermore, eggs from complementation crosses between the *b-4* mutants and the *Bmmah* knockout mutants also exhibited reddish-brown color, which was similar to the *b-4* mutant eggs, indicating that *Bmmah* is responsible for the *b-4* phenotypes.

**Highlight:**

- Responsible region for the *brown egg 4* (*b-4*) mutation was narrowed down by double digest restriction-site associated DNA sequencing (ddRAD-seq).
- The gene structure was disrupted in one of the candidate genes, *Bombyx mori mahogany* (*Bmmah*), in the *b-4* mutant strains.
- CRISPR/Cas9-mediated gene knockout and complementation test confirmed that the *Bmmah* is responsible for the *b-4* phenotypes.
- The *Bmmah* encoded a putative amino acid transporter that belongs to the β-group of solute carrier family.
- The *Bmmah* gene is essential for normal colorization of eggs, compound eyes, and ganglions.

**Graphical abstract:** 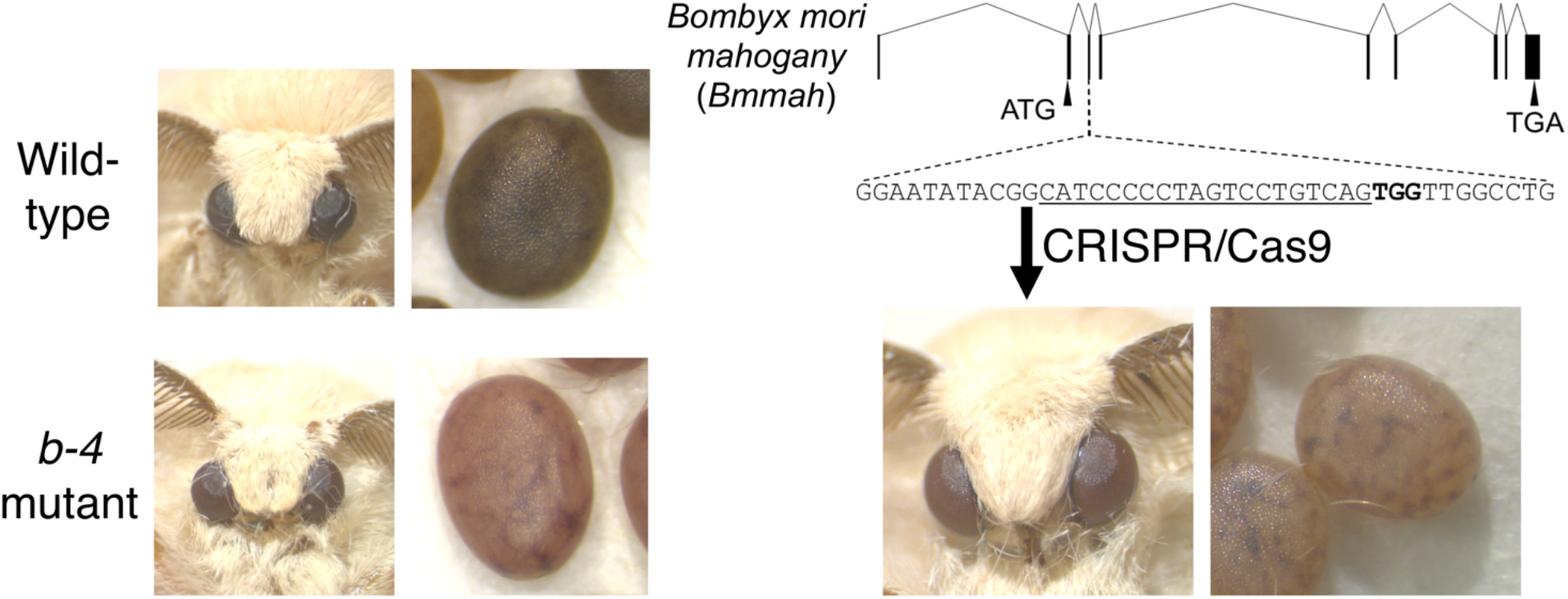

## 1. Introduction

Ommochromes constitute a significant group of pigments that are widely distributed in the eggs, compound eyes, and epidermis of insects (Figon and Casas, 2019). Ommochrome pigments are also known to be involved in body color patterning in invertebrates, which is important for sexual dimorphism and mimicry. For example, *Heliconius* butterflies display mimetic wing patterning, which is regulated by several ommochrome-related genes, such as *vermillion* and *cinnabar* (Reed et al., 2008; Ferguson and Jiggins, 2009). Yellow/red color changes associated with sexual maturation in some dragonflies are regulated by redox states of epidermal ommochrome pigments (Futahashi et al., 2012). In addition, ommochromes also function as screening pigments in compound eyes (Burnet et al., 1968) and are involved in tryptophan metabolism (Linzen, 1974).

Studies on the ommochrome synthetic pathway have been developed by investigating eye color mutants of the fruit fly, *Drosophila melanogaster*. Most of the genes are involved in the synthetic pathway from tryptophan to 3-hydroxykynurenine (*vermillion*: Searles and Voelker, 1986; *cinnabar*: Sullivan et al., 1973), pigment granule formation (*pink*: Falcón-Pérez et al., 2007; *garnet*: Lloyd et al., 1999; *deep orange*: Shestopal et al., 1997), and the incorporation of 3-hydroxykynurenine into pigment granules (*scarlet*: Tearle et al., 1989; *white*: O’Hare et al., 1983). In the silkworm, *Bombyx mori*, ommochrome pigments accumulate in eggs, adult compound eyes, and in the reddish markings on the larval epidermis. The early steps of the ommochrome synthetic pathway are conserved between *B. mori* and *D. melanogaster*. The genes responsible for the three white egg and eye color mutants, *white egg 1* (*w-1*), *white egg 2* (*w-2*), and *white egg 3* (*w-3*), in *B. mori* correspond to the orthologous genes of *cinnabar, scarlet*, and *white* in *D. melanogaster*, respectively (Quan et al., 2007; Kômoto et al., 2009; Tatematsu et al., 2011). The conservation of these genes is also reported in other insects, such as the red flour beetle, *Tribolium castaneum*, and the Western tarnished plant bug, *Lygus hesperus* (Lorenzen et al., 2002; Broehan et al., 2013; Grubbs et al., 2015; Brent and Hull, 2019). However, after 3-hydroxykynurenine incorporation into pigment granules, the ommochrome synthetic pathway begins to differ in *Drosophila* and *Bombyx*. For instance, the gene responsible for the *B. mori red egg* (*re*) mutant encodes a major facilitator superfamily transporter which is conserved in most insect species but lost in *D. melanogaster* (Osanai-Futahashi et al., 2012). This is possibly due to the difference of the ommochrome composition in the ommatidia; while Cyclorrhapha (including *Drosophila*) are reported to have only xanthommatin as the ommochrome pigment in the eye, other insect species, with the exception of Orthoptera, are reported to also contain ommin A in the ommatidia (Linzen, 1974). Given that the *re* gene is indispensable for ommin synthesis (Kawase and Aruga, 1954), the gene loss in Dipteran insects may be the cause of lack of ommins in the compound eyes. Therefore, genetic studies based on non-Cyclorrhapha insect species are important to reveal the later steps of the ommochrome synthesis pathway.

The eggs and adult compound eyes of the *B. mori* contain a mixture of ommochrome pigments, such as ommin and xanthommatin, and exhibit a dark purple or black color (Kawase and Aruga, 1954; Koga and Osanai, 1967; Linzen, 1974; Sawada et al., 2000). The *brown egg 4* (*b-4*) is a recessive mutant which causes eggs and eyes to express a reddish-brown color instead (Fig. 1A; Nakajima, 1956). The *b-4* is located at 21.9 cM in the *B. mori* genetic linkage group 20 (chromosome 20; Doira et al., 1974). It has been reported that although *b-4* mutant and wild-type eggs have similar ommochrome composition, the total amount of pigments are significantly reduced in the mutant eggs (Nakaizawa and Nakajima, 1979). Because the amount of ommochrome precursor, 3-hydroxykynurenine, is comparable between mutant and wild-type eggs, the gene responsible for the *b-4* mutation is predicted to be involved in the synthesis pathway from 3-hydroxykynurenine to ommochrome pigments. We attempted to identify the responsible gene for the mutant phenotypes for a deeper understanding of the later steps of the ommochrome biosynthetic pathway.

**Fig. 1.**
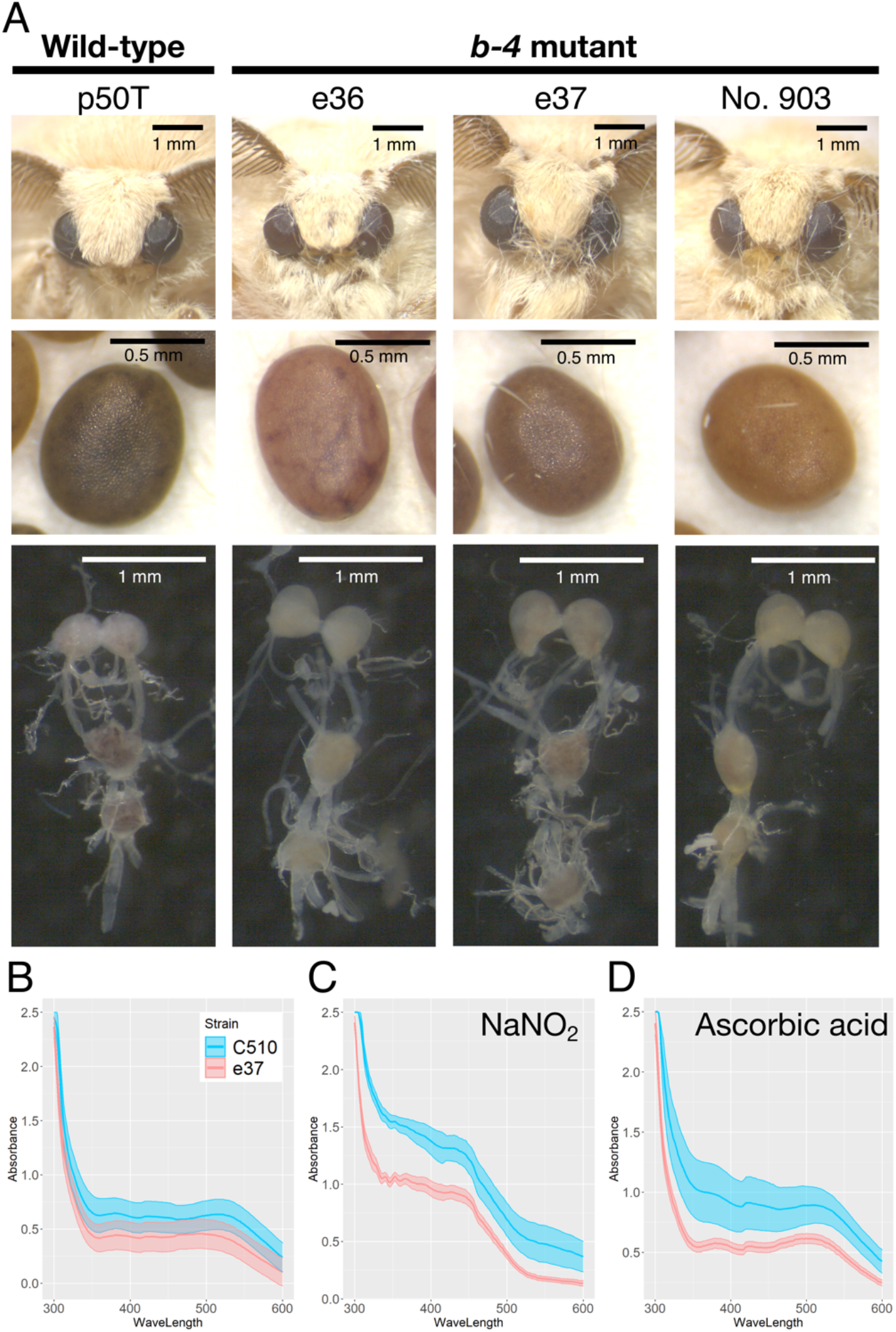
Phenotypes of the *b-4* mutant strains. (A) The observation of the adult compound eyes (top), eggs (middle), and larval brain and ganglions of wild-type (p50T) and *b-4* mutant strains (e36, e37, No. 903). Scale bars are drawn in the images. Group means and standard error bars are presented for the absorption spectrum of egg pigments in (B) normal state, (C) oxidized state, and (D) reduced state.

Here, we performed double digest restriction-site associated DNA sequencing (ddRAD-seq) and narrowed down the *b-4* locus to approximately 1.1 Mb that contains 69 predicted gene models. Among these, three *b-4* mutant strains investigated in this study had two independent mutations in an ortholog of the *Drosophila* eye color mutant gene, *mahogany* (*mah*), which encodes a putative amino acid transporter that belongs to the β-group of solute carrier (SLC) family (Grant et al., 2016). We performed CRISPR/Cas9-mediated gene knockout targeting the candidate gene and confirmed that the responsible gene for the *b-4* phenotypes is *B*. *mori mahogany* (*Bmmah*).

## 2. Materials and Methods

### 2.1. B. mori strains

We used three wild-type *B. mori* strains (p50T, C108, and C510) and three *b-4* mutant strains (e36, e37, and No. 903) for this study. The *B. mori* strain, p50T, was maintained in our laboratory. Strains e36 and e37 were provided from Kyushu University (Fukuoka, Japan) with the support from the National Bioresource Project (http://silkworm.nbrp.jp), and C108, C510, and No. 903 were acquired from the Genetic Resource Center, National Agriculture and Food Research Organization (NARO). *B. mori* larvae were reared on fresh mulberry leaves or an artificial diet (SilkMate PS (NOSAN)) under continuous 12 h-light/12 h-dark conditions at 25 °C.

### 2.2. Wave scanning of pigments by spectrophotometer

Ten eggs were prepared from 3 batches of strains C510 and e37. The eggs were homogenized by plastic pestle in 200 μL of 2% hydrochloric acid in methanol (v/v) to extract pigments. We added an oxidizing agent (1% NaNO_2_) and a reducing agent (1% ascorbic acid) into every extract to observe redox-dependent color changes. After centrifugation (3000 × g for 5 min), supernatants were collected and used for analyses. Wave scanning of samples was performed using a GeneQuant 1300 spectrophotometer (Biochrom).

### 2.3. ddRAD-seq and mapping of the b-4 locus

We obtained F_1_ individuals from a single pair cross between the *b-4* mutant (e36) female and the wild-type (p50T) male. We then crossed F_1_ moths with each other and obtained F_2_ eggs. Genomic DNAs were extracted from each F_2_ neonate larva (31 reddish-brown eggs and 61 normally pigmented eggs), F_1_ moths, and parental moths, using the Maxwell 16 Tissue DNA Purification kit (Promega). The ddRAD-seq library was constructed using all the genomic DNAs according to the method described in Peterson et al. (2012). Genomic DNAs from these individuals were double digested with EcoRI and MspI and served for making ddRAD-seq libraries. A unique pair of inline barcodes was ligated onto the EcoRI-associated DNA and MspI-associated DNA for each sample to identify each variety. The ddRAD-seq library was then sequenced on one lane using an Illumina HiSeq 2500, generating 200-bp paired-end reads. Only the sequenced reads in which the beginning of the forward and reverse reads matched the barcode 100% were extracted. After removing the barcodes, the reads were demultiplexed by sample. The demultiplexed reads were then mapped to the *B. mori* genome (Kawamoto et al., 2019) that was downloaded from the *B. mori* genome database (SilkBase; http://silkbase.ab.a.u-tokyo.ac.jp/) by BWA mem (ver 0.7.15) (Li, 2013). We extracted the SNPs using the UnifiedGenotype module of the Genome Analysis Toolkit (ver 3.3-0) (McKenna et al., 2010; DePristo et al., 2011). Genotypic biases between individuals with wild-type and mutant phenotypes were tested using a Fisher’s exact test, and a Manhattan plot was drawn using linkage analysis~manhattan plot module in Portable Pipeline (https://github.com/c2997108/OpenPortablePipeline). The *b-4* locus was narrowed down by Excel so that the phenotype and genotype matched 100%.

### 2.4. RNA-sequencing (RNA-seq)

Eggs of the *b-4* mutant strain, e37, were collected at five time points (0, 24, 48, 72, and 120 hours post-oviposition (hpo)). Total RNA was extracted from the egg samples using ISOGEN (NIPPON GENE) and purified with the RNeasy Mini Kit (QIAGEN). RNA-seq libraries were prepared using the TruSeq Sample Preparation Kit v2 (Illumina) and sequenced on a HiSeq 2500 platform (Illumina) with 200 bp paired-end reads. RNA-seq reads of wild-type strains (C108 and p50T) were obtained from DDBJ accession number DRA003068 (Osanai-Futahashi et al., 2016). Low-quality reads and adapter sequences were removed using Trim Galore! (https://github.com/FelixKrueger/TrimGalore/issues/25). We mapped the RNA-seq reads to the *B. mori* genome (Kawamoto et al., 2019) with HISAT2 (Kim et al., 2019). Generated Binary Alignment Map (BAM) files were loaded into ggsashimi (Garrido-Martín et al., 2018) to visualize splicing variants with Sashimi plots. The transcript expression was quantified using StringTie (Pertea et al., 2015) and differentially expressed genes were identified using edgeR (Robinson et al., 2010). We assembled RNA-seq reads using Trinity (Grabherr et al., 2011) to determine deduced mRNA sequences in the mutant strain, e37.

### 2.5. Reverse transcription-polymerase chain reaction (RT-PCR), genomic PCR, and sequencing

Total RNA was isolated from the eggs at 24 hpo from a wild-type strain (p50T) and three *b-4* mutant strains (e36, e37, and No. 903) using TRIzol (Invitrogen). Reverse transcription was then performed using the oligo (dT) primer and Avian myeloblastosis virus reverse transcriptase contained in the TaKaRa RNA PCR kit (TaKaRa). We conducted RT-PCR using KOD One (TOYOBO) with the primer sets listed in Table S1 under the following conditions: initial denaturation at 94 °C for 1 min, 45 cycles of denaturation at 98 °C for 10 sec, annealing at 60 °C for 5 sec, and extension at 68 °C for 10 sec. Genomic DNA was subsequently extracted from adult legs with the DNeasy Blood & Tissue Kit (QIAGEN) according to the manufacturer’s protocol. Genomic PCR was conducted using KOD One with the primer sets listed in Table S1 under the following conditions: 94 °C for 1 min; 30 cycles of 98°C for 10 sec, 60–62°C for 5 sec, and 68°C for 5 sec/kb. The PCR products were then sequenced using the ABI3130xl genetic analyzer (Applied Biosystems). Transmembrane prediction was performed using TOPCONS (Tsirigos et al., 2015). The graphical images of the transmembrane proteins were generated using the PROTTER program (Omasits et al., 2014).

### 2.6. CRISPR/Cas9-mediated gene knockout

Unique CRISPR-RNA (crRNA) target sequences in the *B. mori* genome were selected using CRISPRdirect (https://crispr.dbcls.jp/; Naito et al., 2015). A mixture of crRNA (200 ng/μL; FASMAC), trans-activating crRNA (tracrRNA) (200 ng/μL; FASMAC), and Cas9 Nuclease protein NLS (300 ng/μL; NIPPON GENE) in injection buffer (100 mM KOAc, 2 mM Mg(OAc)_2_, 30 mM HEPES-KOH; pH 7.4), was injected into each egg within 3 h after oviposition (Yamaguchi et al., 2011). The injected embryos were then incubated at 25 °C in a humidified Petri dish until hatching.

Adult generation 0 (G_0_) moths were crossed with wild-type (p50T) moths to obtain generation 1 (G_1_). Genomic DNA was extracted from the knockout G_1_ individuals by the HotSHOT method (Truett et al., 2000), and Genomic PCR was performed as described above. Mutations at the targeted site were detected by heteroduplex mobility assay using the MultiNA microchip electrophoresis system (SHIMADZU) with the DNA-500 reagent kit (Ota et al., 2013; Ansai et al., 2014). G_1_ individuals carrying a heterozygous knockout mutation were intercrossed with each other to generate homozygous knockout G_2_ individuals. The PCR products of the targeted site in G_2_ were used for DNA sequencing. The G_2_ individuals carrying a homozygous knockout mutation were crossed with each other to establish knockout strains.

### 2.7. Phylogenetic analysis

#### 2.7.1 Phylogenetic analysis of KWMTBOMO12119 homologous proteins in insects

We collected proteins homologous to BmMAH in the NCBI RefSeq protein database (Lepidoptera; *B. mori, Trichoplusia ni*, Diptera; *D. melanogaster, Musca domestica, Aedes aegypti*, Hymenoptera; *Apis mellifera, Bombus terrestris*, Coleoptera; *T. castaneum, Nicrophorus vespilloides*, Hemiptera; *Acyrthosiphon pisum, Cimex lectularius*, Blattodea; *Zootermopsis nevadensis*) using the BLASTP program at an e-value cutoff of 1e-10. All sequences were aligned using the MUSCLE program (Edgar, 2004), and a phylogenetic tree was constructed by MEGA X software (Kumar et al., 2018) using the maximum likelihood method with 1000 bootstrap replications. All positions with less than 80% site coverage were eliminated.

#### 2.7.2 Phylogenetic analysis of proteins relative to SLC32, 36, and SLC38

The human protein sequences for the β-group of SLC families were obtained from the Ensembl all-peptide datasets (http://m.ensembl.org/) according to the annotations. Protein sequences for the β-group of SLC families in *Mus musculus, Danio rerio, D. melanogaster, Caenorhabditis elegans*, and *B. mori* were collected from the Ensembl all-peptide datasets or SilkBase protein dataset. For each protein dataset, proteins that had a consensus motif of the β-group of SLCs (*i.e*. Pfam “Aa_trans” motif; https://pfam.xfam.org/family/Aa_trans) were searched using HMMsearch (Eddy, 2009). All hit proteins with significance (E-value < 10) in HMMserch were filtered against the human RefSeq protein database using the BLASTP program. Only those that hit a sequence from human SLC32, 36, and 38 families with the highest score were used for the phylogenetic analysis. The protein alignment and the phylogenetic tree construction were performed as described above.

## 3. Results

### 3.1. Characterization of the b-4 phenotype

Eggs and adult compound eyes of the wild-type strain, p50T, exhibited a dark purple or black color, while that of the *b-4* mutant strains e36, e37, and No. 903 displayed a dark reddish-brown color (Fig. 1A). However, the phenotypes were slightly different between the *b-4* mutant strains. Compared to e36 and No. 903, the e37 strain had darker eggs and compound eyes. We also observed a reduction of pigments in the brain and ganglion cells of the *b-4* mutants (Fig. 1A), which is unreported in previous research (Nakajima, 1956; Nakaizawa and Nakajima, 1979).

To analyze the composition of egg pigments between the wild-type strain, C510, and the *b-4* mutant strain, e37, we extracted the pigments with acidic methanol. We then measured the spectrum of the pigments using a spectrophotometer with or without an oxidizing (NaNO_2_) or a reducing (ascorbic acid) agent to observe redox-dependent color changes (Fig. 1B-D). Although the total amount of pigments were reduced in the mutant eggs, there was no significant difference in the waveform regardless of redox conditions. We obtained similar results when using pigments extracted from the brains and ganglion cells (Fig. S1).

### 3.2. Narrowing down the b-4 locus by ddRAD-seq

We performed genetic linkage analysis using the ddRAD-seq to identify the candidate region linked to the *b-4*. First, we obtained F_1_ individuals by crossing a single female of a *b-4* mutant strain, e36, with a single male of a wild-type strain, p50T. We crossed F_1_ moths with each other and obtained F_2_ eggs. Genomic DNA extracted from 92 F_2_ individuals (31 reddish-brown eggs, 61 normally pigmented eggs), a single pair of F_1_ hybrids, and a single pair of parental moths were subjected to the ddRAD-seq analysis. Based on the single nucleotide polymorphisms (SNPs) data obtained from the analysis, we narrowed down the *b-4*-linked region to approximately 1.1 Mb on chromosome 20, from position 778674 to 8897242 (Fig. 2). The region contained 69 predicted gene models according to the recent *B. mori* gene list (Kawamoto et al., 2019) (Table S2).

**Fig. 2.**
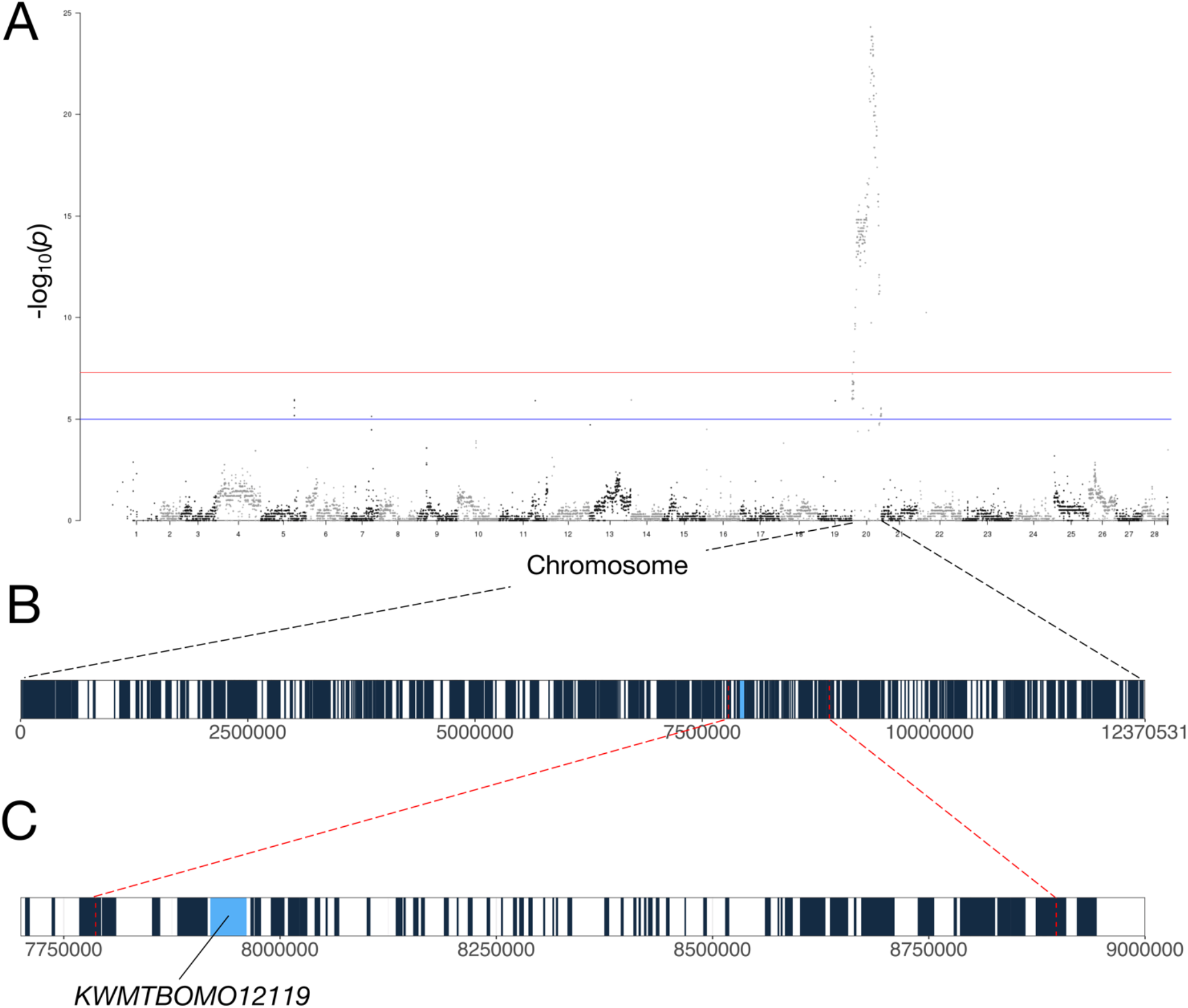
Mapping of the *b-4* candidate region in chromosome 20. (A) Manhattan plot for ddRAD-seq analysis in F_2_ individuals. The blue line and red line indicate p-value = 1.0×10^-5^ and 5.0×10^-8^, respectively. A significant (p-value < 5.0×10^-8^) peak was observed only on chromosome 20. Predicted gene models on (B) chromosome 20 and (C) the *b-4* candidate region. The red dotted lines indicate borders of the responsible region for the *b-4* locus narrowed by ddRAD-seq. Each filled rectangle represents a gene model. The *b-4* candidate gene *KWMTBOMO12119* is colored in blue.

### 3.3. Transcript analysis of the b-4 candidate genes

Color differences between *b-4* and wild-type eggs become distinct 48–72 hpo. According to our previous research, the genes responsible for eye color mutants, such as *Bm-white* (*Bmwh3*), *Bm-vermillion, Bm-cinnabar* (*BmKMO*) *Bm-re, Bm-scarlet* (*Bm-w-2*), and *Bm-cardinal*, are all highly expressed in 24–48 hours after the eggs are laid (Osanai-Futahashi et al., 2016). To observe mRNA expression of all 69 candidate gene models, we calculated fragments per kilobase of exon per million reads mapped (FPKM) using previous RNA-seq data from two wild-type strains, p50T and C108, at four time points (0, 24, 48, and 72 hpo, Table S2). Among the candidates, 18 genes were highly expressed (FPKM> 10) in either 24 or 48 hpo in both strains. Of these, we only focused on 5 gene models (*KWMTBOMO12119, 12123, 12124, 12125, 12178*) that were zygotically expressed in 24 hpo because the *b-4* egg color phenotype is not subject to maternal inheritance, and transcripts of the other 13 candidate genes were detected in eggs just after oviposition (Table S2). Furthermore, we noticed that *KWMTBOMO12123, 12124*, and *12125* code a single gene that we collectively named *KWMTBOMO12123-5*. Therefore, 3 genes were possible candidates for the cause of the *b-4* phenotype.

We performed RNA-seq analysis using eggs of a mutant strain, e37, to compare the expression pattern and deduce amino acid sequences of the candidate genes with wild-type strains. None of the 3 candidates were differentially expressed (False Discovery Rate; FDR> 0.01) in 24–48 hpo between strains p50T and e37 (Table S3). *De novo* assembled transcript sequences revealed that one and two amino acids were substituted in genes *KWMTBOMO12119* and *KWMTBOMO12123-5*, respectively, while there were no substitutions in gene *KWMTBOMO12178* between the two strains (Table S3). Every substitution found in *KWMTBOMO12119* and *KWMTBOMO12123-5* was also observed in *+^b-4^* strains according to the hypothetically reconstructed genome data in SilkBase (Table S3). We then viewed the mapped reads on these three gene models and noticed that a major transcript of *KWMTBOMO12119* began from exon 5 in the *b-4* strain, e37 (Fig. 3A, Fig. S2), suggesting that most proteins translated from this gene are functionally disrupted. The *KWMTBOMO12119* codes a putative amino acid transporter which has 11 transmembrane domains (Fig. 3). However, proteins translated from the major transcript of the *b-4* e37 strain were predicted to have only seven transmembrane domains (Fig. 3), suggesting that the abnormal transcription of *KWMTBOMO12119* is the cause of the *b-4* phenotype.

**Fig. 3.**
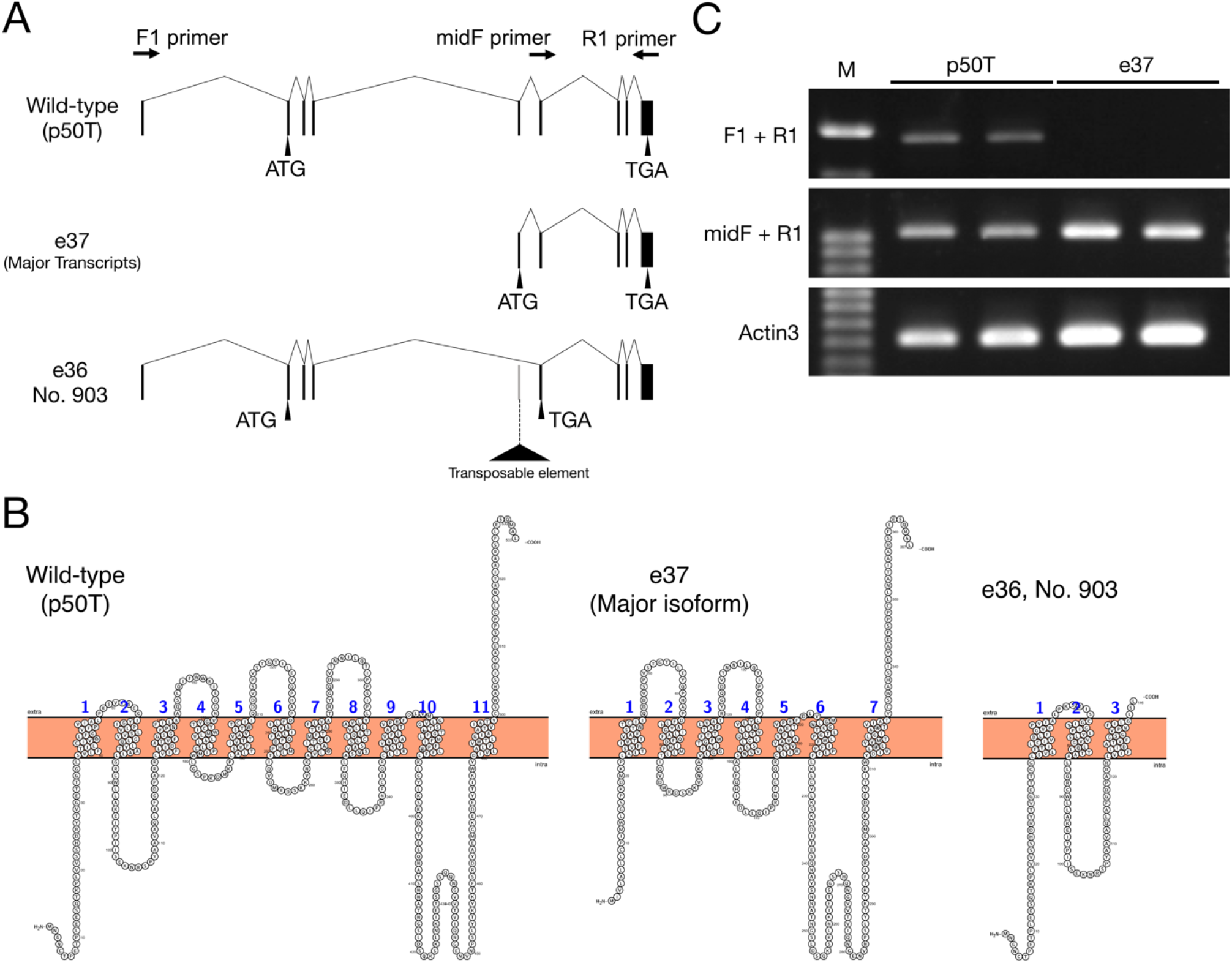
Gene structure of the *b-4* candidate gene *KWMTBOMO12119*. (A) Exon structures of the *KWMTBOMO12119* gene in wild-type (p50T) and *b-4* mutant strains (e37, e36, No. 903). The *KWMTBOMO12119* is mainly transcribed from the exon 5 in a *b-4* mutant strain e37. The exon 5 is skipped in *b-4* mutant strains, e36 and No. 903, due to a transposable element insertion. Arrows indicate the position of primers, Bmmah_CDS_F1 (F_1_), Bmmah_CDS_midF (midF), and Bmmah_CDS_R1 (R1) listed in Table S1, respectively. (B) The transmembrane structure of KWMTBOMO12119 protein in the wild-type strain, p50T, and *b-4* mutant strains, e37, e36 and No. 903. (C) Gel electrophoresis of RT-PCR products amplified from cDNA obtained from eggs of wild-type (p50T) and e37. Primer sets used for RT-PCR are shown in left. The *BmActin3* gene are analyzed as a control.

### 3.4. *Sequencing analysis of the b-4 candidate gene KWMTBOMO12119* in *b-4* mutants

We isolated total RNA from the eggs at 24 hpo of the *b-4* (e36, e37, No. 903) and wild-type (p50T) individuals. Each cDNA was obtained by reverse transcription. Sequencing of the cDNAs revealed that exon 5 is skipped in two of three mutant strains, e36 and No. 903. Genomic PCR found an approximately 4.8 kb insertion at the boundary between exon 5 and intron 5 (Fig. 3A, Fig. S3). A BLAST search of the inserted DNA sequence showed significant homology to a *B. mori* transposable element, SINE:Bm1 (Adams et al., 1986). The exon skipping causes a frame-shift and premature stop codon, resulting in the production of truncated proteins (Fig. 3). RT-PCR confirmed that major transcripts of *KWMTBOMO12119* were initiated from exon 5 in the e37 strain (Fig. 3B). Furthermore, intact coding sequence (CDS) of *KWMTBOMO12119* was also obtained from the e37 strain cDNA (data not shown), indicating that functional *KWMTBOMO12119* transcripts, including exon 1–4, exist in the mutant strain. This is consistent with the RNA-seq results and phenotypes of the darker egg and compound eye colors in the e37 strain compared to those in the other two *b-4* strains (Fig. 1A).

## 3.5. Functional analysis of the KWMTBOMO12119 gene by CRISPR/Cas9-mediated gene knockout

To investigate whether the *KWMTBOMO12119* gene is essential for normal egg and eye pigmentation, we introduced mutations into the candidate gene by CRISPR/Cas9 system. We selected a specific target site in exon 3 of the *KWMTBOMO12119* gene (Fig. 4A). In the G_0_ generation, 18 of the 24 adult moths obtained possessed partially or entirely reddish-brown compound eyes (Fig. 4B, Table S4). Using some G_0_ individuals, we established three *KWMTBOMO12119* knockout strains, and sequence analysis showed that a single different mutation was induced in each strain (Fig. 4C). The strains carrying a four-nucleotide deletion, a five-nucleotide deletion, and a nine-nucleotide deletion were termed *Δ4, Δ5*, and *Δ9*, respectively. The former two alleles result in frame-shifts and premature stop codons, whereas the latter results in three amino acid deletions in the second transmembrane region. All three strains possessed reddish-brown eggs and adult compound eyes (Fig. 4D). We also observed a reduction of pigments in the brain and ganglion cells of the *KWMTBOMO12119* knockout mutants (Fig. S4).

**Fig. 4.**
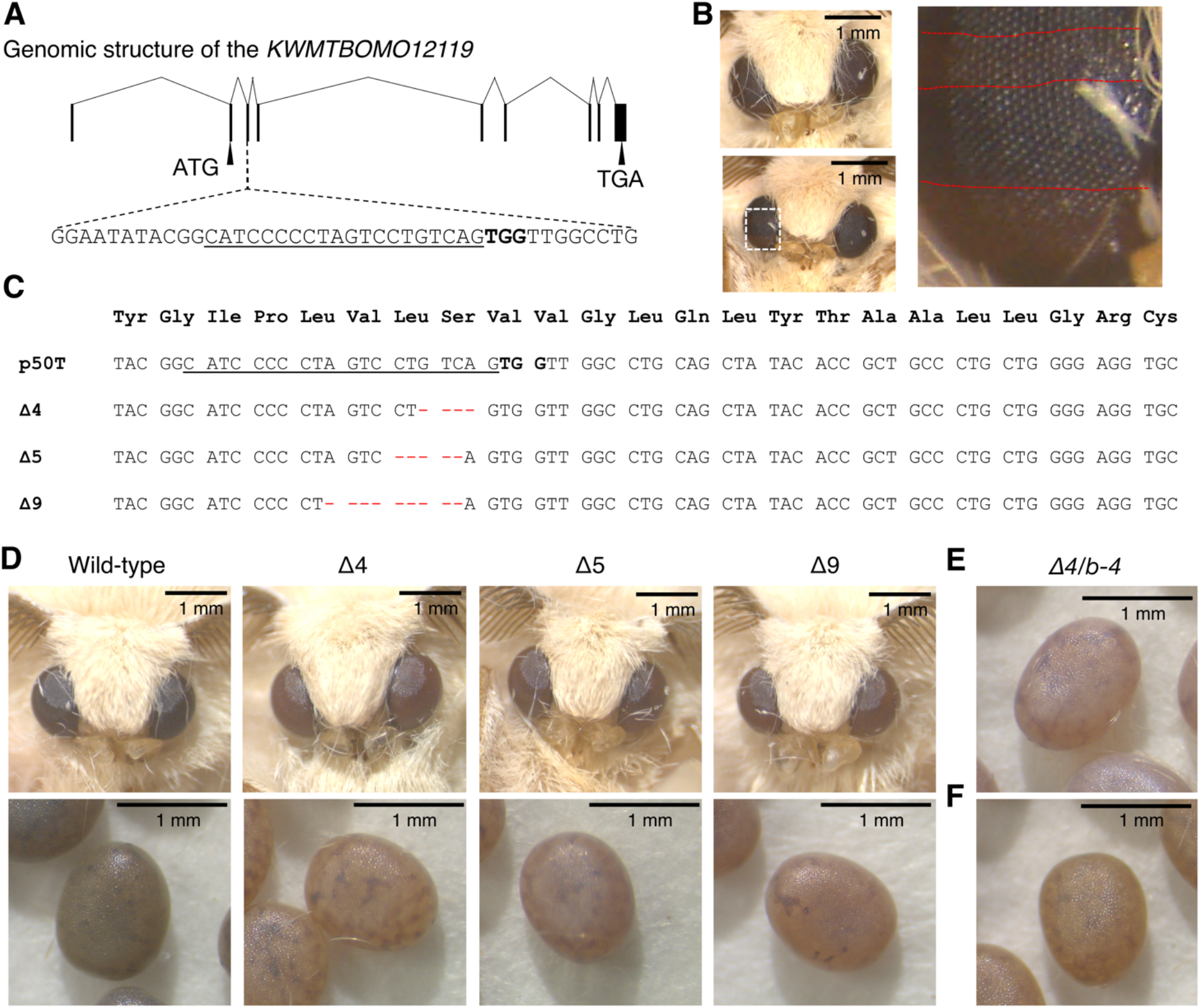
CRISPR/Cas9-mediated knockout of the *KWMTBOMO12119* gene. (A) The exon structure of the *KWMTBOMO12119* gene and a crRNA target site. The target sequence is underlined and the protospacer adjacent motif (PAM) is indicated by bold letters. (B) Representative phenotype of G_0_ adult moths that retain entirely (upper left) or partially (lower left) reddish-brown compound eyes. An enlarged view of the partially reddish-brown compound eye is shown to the right. The red dotted lines outline the reddish-brown colored area. Scale bars: 1 mm. (C) Alignment of the *KWMTBOMO12119* gene sequences surrounding the crRNA target site from wild types (p50T), and G_2_ individuals with a single *KWMTBOMO12119* knockout allele (*Δ4, Δ5, Δ9*). The target sequence is indicated by underline and the PAM is written in bold letters. (D) Representative reddish-brown compound eyes of an adult *KWMTBOMO12119* knockout moths (top) and eggs (bottom). Scale bar: 1 mm. (E-F) F_1_ eggs obtained from a complementary cross (E) between e37 female and *Δ4* strain male, and (F) between *Δ4* strain female and e37 male. Scale bar: 1mm.

In complementary crosses between an e36 female and *Δ4* strain male, F_1_ eggs exhibited reddish-brown color (Fig. 4E) that is similar to the *b-4* mutant phenotype. This similar phenotype was observed in reciprocal mating between a *Δ4* strain female and e36 male (Fig. 4F). These results indicate that the mutations in the gene are responsible for the *b-4* egg color phenotype.

## 3.6. Phylogenetic analysis of KWMTBOMO12119 protein

We collected sequences of proteins homologous to KWMTBOMO12119 from 12 insects whose whole genome sequences have been published and constructed a phylogenetic tree (Fig. 5A). The phylogenetic tree showed that *KWMTBOMO12119* is orthologous to the *D. melanogaster* eye color mutant gene, *mahogany* (*mah*). Thus, we designated KWMTBOMO12119 as BmMAH. We further found that *B*. *mori* sex-linked translucent (OS) protein, which is known to be involved in uric acid accumulation in the larval integument, is a sister group of BmMAH. As we reported previously, the OS orthologs are poorly conserved among insects (Kiuchi et al., 2011; Tomihara et al., 2019), whereas the BmMAH orthologs are widely conserved in insects, except for the termite *Z. nevadensis*. Insect species that lack both BmMAH and OS orthologs could not be found in the scope of the species we searched. In addition, BmMAH and OS have high homology with mammalian SLC32, 36, and 38, which are together clustered into the β-group of SLCs (Schiöth et al., 2013). Thus, we performed a phylogenetic analysis using protein sequences relative to the β-group of SLCs in mammals (*H. sapience* and *M. musculus*), fish (*D. rerio*), nematodes (*C. elegans*), and insects (*D. melanogaster* and *B. mori*) (Fig. 5B). The results confirmed that BmMAH and OS belong to the β-group of SLCs, although the proteins′ closest relatives were not conserved in vertebrates.

**Fig. 5.**
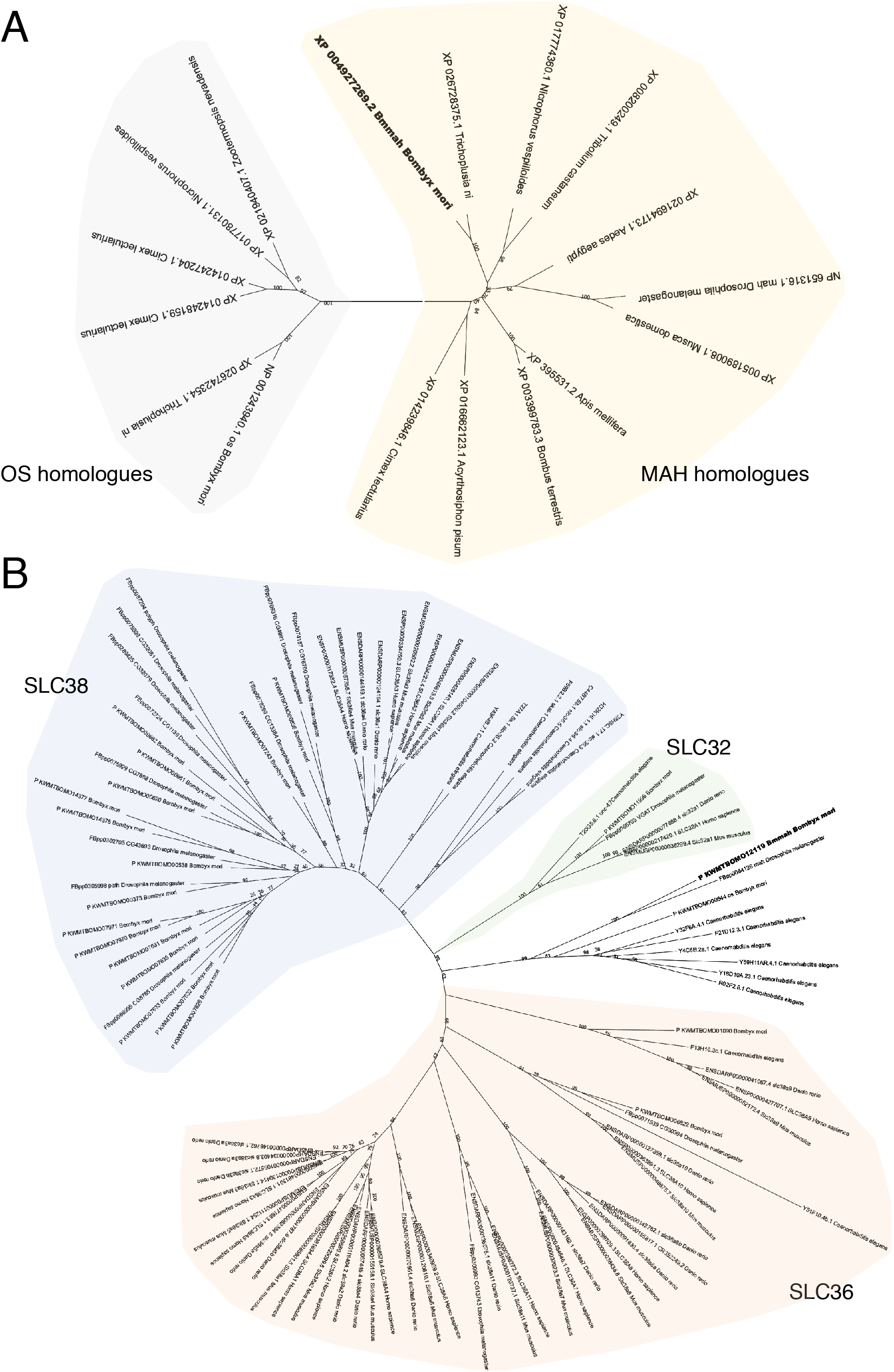
Phylogenetic analysis of KWMTBOMO12119 homologous proteins. (A) Phylogenetic tree of KWMTBOMO12119 homologous proteins in insects. NCBI accession numbers are given with species names. (B) Phylogenetic tree of proteins belongs to β-group of SLCs in mammals, fish, nematode, and insects. Ensembl accession numbers and *B. mori* gene model names are given with species names. KWMTBOMO12119 protein (XP004927269.1, BmMAH) is shown in bold letters.

## 4. Discussion

In this research, we performed ddRAD-seq analysis and narrowed down the *b-4* candidate region to a segment that contained 69 predicted gene models (Fig. 2). RNA-seq and cDNA sequencing revealed that the gene structure is disrupted in one of the candidate genes, *KWMTBOMO12119*, in the *b-4* mutant strains (Fig. 3, Fig. S2-3). Next, we performed CRISPR/Cas9 mediated gene knockout targeting *KWMTBOMO12119* and observed reddish-brown colored eggs and adult compound eyes like in the *b-4* mutants (Fig. 4B, D). In addition, F_1_ eggs obtained from complementary crosses between the *b-4* mutant and the gene knockout moths also exhibited a reddish-brown color similar to that of the *b-4* mutants (Fig. 4E-F), demonstrating that the mutations in *KWMTBOMO12119* are responsible for the *b-4* phenotypes. The phylogenetic analysis showed that *KWMTBOMO12119* is the ortholog to an eye color gene in *D. melanogaster, mah*, which encodes an SLC transporter (Beadle and Ephrussi, 1937; Grant et al., 2016) (Fig. 5). Accordingly, we named the *KWMTBOMO12119* gene *B. mori mah* (*Bmmah*).

Nakaizawa and Nakajima (1979) reported that *b-4* mutant and wild-type eggs have similar ommochrome compositions, while the total amount of the pigments are significantly reduced in the mutant eggs. The ommochrome precursor, 3-hydroxykynurenine, accumulates in the mother’s pupal ovaries (Yamashita and Hasegawa, 1964) and is incorporated into pigment granules in eggs by the heterodimeric ATP-binding cassette (ABC) transporters encoded by *Bm-w-2* and *Bmwh3* after oviposition (Tatematsu et al., 2011). The *b-4* egg color phenotype is not subject to maternal inheritance like in the egg color mutants, *w-2, w-3, pe* (*pink-eyed white egg*), and *re*. Furthermore, about three-quarters of G_2_ eggs obtained from intercrosses between G_1_ individuals (i.e. heterozygous mutants) exhibited normal pigmentation (data not shown), suggesting that the amount of 3-hydroxykynurenine in the *b-4* mutant eggs is comparable to that in wild-type. An increased expression of *Bmmah* mRNA was observed at 24 hpo, just before egg pigmentation occurred (Table S2). This is similar to other egg color mutant genes, such as *Bm-w-2, Bmwh3, Bm-cardinal*, and *Bm-re* (Osanai-Futahashi et al., 2016), all of which are involved in the latter step of the ommochrome biosynthesis pathway after 3-hydroxykynurenine is incorporated into eggs (Kômoto et al., 2009; Tatematsu et al., 2011; Osanai-Futahashi et al., 2012, 2016). These results suggest that the BmMAH transporter either participates in the transportation of 3-hydroxykynurenine into pigment granules or the conversion of 3-hydroxykynurenine to final ommochromes by the transportation of unknown substances. However, since the ABC transporters, *white* and *scarlet*, are reported to localize in pigment granules in the *Drosophila* ommatidia (Mackenzie et al., 2000) and their corresponding *B. mori* mutants have white eggs, it is more likely that BmMAH transports undefined substances that are directly or indirectly involved in the conversion of 3-hydroxykynurenine to final ommochromes.

The phylogenetic tree revealed that the BmMAH is a paralog of the OS protein (Fig. 5) which is involved in uric acid accumulation in the larval integument (Tamura and Akai, 1990; Kiuchi et al., 2011; Tomihara et al., 2019). BmMAH appears to be highly conserved among insects, including hemimetabola, such as the pea aphid, *A. pisum*, and the bed bug, *C. lectularius*, although *Z. nevadensis*, was found to only possess OS (Fig. 5). Other insect species, such as the western honey bee, *A*. *mellifera, D. melanogaster*, and *T. castaneum*, have lost OS, while insects that lack both BmMAH and OS could not be found in this study, suggesting that BmMAH and OS share some redundant functions. In fact, egg colors of the *os* mutants and the *os* knockout mutants are slightly lighter than the wild-type, suggesting that OS is also involved in the ommochrome pigmentation (Fig. S5; Kiuchi et al., 2011). On the other hand, the integument color of *Bmmah* mutants does not show traces of transparency (Fig. S6) and the eye color of *os* mutants are normal. This can be explained by the expression pattern of these transporter genes. The mRNA level of the *Bmmah* gene is about one-eighth of that of the *os* gene in the integument but comparable in the embryo according to our previous RNA-seq data (SilkBase assembled RNA-seq libraries A_BomoSK and A_BomoEE).

In the ommochrome pigment granules of *D. melanogaster, B. mori*, and *T. castaneum*, the oxidative condensation of 3-hydroxykynurenine to xanthommatin is supposed to be catalyzed by a heme peroxidase, Cardinal, although biochemical data are not sufficient (Howells et al., 1977; Harris et al., 2011; Osanai-Futahashi et al., 2016). Xanthine dehydrogenase (XDH) is incorporated into pigment granules and probably involved in pteridine biosynthesis in *D. melanogaster* (Reaume et al., 1989, 1991). In addition, recent research indicates that uric acid is synthesized by XDH from its precursors, xanthine and hypoxanthine, in urate granules of *B. mori* integument (Fujii and Banno, 2019). Thus, it is possible that BmMAH and OS transport a substance commonly involved in the biosynthesis of ommochromes, pteridines, and uric acid.

Ommochrome pigment granules “ommochromasomes” and urate granules are classified as a group of cell type-specific compartments known as lysosome-related organelles (LROs), such as mammalian melanosomes. In *D. melanogaster* eye color mutants, both ommochrome and pteridine pigments are reduced by mutations in most genes involved in LRO biogenesis, such as *carmine, garnet*, and *pink* (Ferre et al., 1986). The *B. mori distinct oily* (*od*) mutant exhibits a reduction of uric acid and ommochromes in the larval integument and eggs, respectively (Hatamura, 1939; Fujii et al., 2008). The gene responsible for the *od* phenotypes is a *B. mori* homolog of the human *biogenesis of LRO complex1, subunit 2* (*BmBLOS2*) Although most properties of LROs are shared with lysosomes, these organelles contain cell type-specific components that are responsible for their specialized functions (Dell’Angelica et al., 2000). For example, tyrosinase and tyrosinase-related protein 1 are transported to the melanosomes and contribute to melanin synthesis in melanocytes of skin and eyes in mammals (Wasmeier et al., 2008).

The closest homologs of BmMAH and OS in mammals are the β-group of SLCs, which includes SLC32, 36, and 38 (Fig. 5B). It is reported that some of the β-group of SLCs transport amino acids (Schiöth et al., 2013), most of which are neither ommochrome nor uric acid precursors. The only member of the SLC32 family in mammals, SLC32A1, mediates H^+^ driven uptake of the inhibitory amino acids, γ-aminobutyric acid (GABA) and glycine, into synaptic vesicles (McIntire et al., 1997; Chaudhry et al., 1998; Farsi et al., 2016), and GABAergic synaptic vesicles have higher pH (~6.4) than glutamatergic synaptic vesicles (pH ~5.8) (Egashira et al., 2016). Moreover, in SLC32A1-deficient neurons, the pH in GABAergic synaptic vesicles is significantly reduced (Egashira et al., 2016). One of the SLC36 transporters, SLC36A1, localizes to lysosomes and cotransports neutral amino acids with H^+^ (Sagné et al., 2001). Some SLC38 transporters, such as SLC38A7 and SLC38A9, localize to lysosomal membranes and function as Na^+^-dependent amino acid transporters (Verdon et al., 2017). Na^+^ uptake into the lysosomal lumen can hinder the acidification because membrane potential is determined by concentration gradients of all ions between the lumen and cytosol (Xiong and Zhu, 2016). Notably, a two-pore sodium channel (TPC2), a chloride channel (oculocutaneous albinism type 2; OCA2), and the SLC45A2 transporter regulate melanosome pH to support melanization (Bellono et al., 2016; Le et al., 2020). From these observations, we speculate that BmMAH transports some amino acids on the membrane of pigment granules and maintains the luminal pH, which contributes to normal ommochrome synthesis. Although quantification of the ommochromasome pH has never been performed, other LROs share characteristics with lysosomes, such as acidic pH (Figon and Casas, 2019). An abnormal pH of eye pigment granules may also inhibit pteridine synthesis in *D. melanogaster*. This may be the reason why pteridine pigments are also reduced in the pigment granules of the *mah* mutant (Ferre et al., 1986). The same scenario can explain the reduction of uric acid in the integument of *os* mutants where there is inhibition of uric acid synthesis under abnormal pH condition of urate granules. Intriguingly, a missense mutation in *SLC36A1* in horses results in a reduction of pigmentation in the eyes, skin, and hair (Cook et al., 2008), indicating that some β-group of SLCs contribute to a variety of pigment systems in a broad range of animals. An investigation into the localization of BmMAH and OS at the membrane of pigment and urate granules, as well as the substrate of these transporters, is required to elucidate our hypothesis.

## Acknowledgements

We are grateful to Natsuki Nakashima for technical assistance for silkworm maintenance. We also thank Seigo Kuwazaki for ddRADseq library preparation. We thank the Institute for Sustainable Agro-ecosystem Services, The University of Tokyo, for facilitating the mulberry cultivation and the Biotron Facility at the University of Tokyo for rearing the silkworms. This work was supported by JSPS KAKENHI (grant numbers 17H05047 and 20H02997) and grant from the Ministry of Agriculture, Forestry and Fisheries of Japan (Research Project for Sericultural Revolution) to TK. This work was also supported by JSPS KAKENHI Grant Number JP20J22954 to KT, JSPS KAKENHI Grant Number 26850220 and Ibaraki University Grant for Specially Promoted Research to M. O-F, and Cooperative Research Grant of the Genome Research for BioResource, NODAI Genome Research Center, Tokyo University of Agriculture, and the National BioResource Project, Japan.

## Data Availability

RNA-seq data was submitted to DDBJ under accession numbers DRA011661-DRA011663, DRA011665-DRA011666, DRA011668-DRA011672. The nucleotide sequences of *Bmmah* were submitted to the DDBJ under accession numbers of LC616418 (p50T), LC616419 (e37 minor isoform), LC617101 (e37 major isoform), and LC616420 (e36 and No.903).

## Author contributions

KT, MO-F and TK designed the study. KT conducted most of the experiments. KS and TK helped CRISPR/Cas9-mediated gene knockout and SM and MO-F performed wave scanning of the pigments. MO-F reared and crossed the silkworms for ddRAD-seq analysis and embryonic RNA-seq analysis. MO-F and K Yamamoto prepared the RAD-seq library, and RF prepared the RNA-seq library. HU and SY performed Hiseq2500 sequencing experiments. K Yoshitake, MO-F, RF, and KT analyzed the sequence data. KT and TK wrote the manuscript, with input from SK, K Yoshitake and MO-F. All authors approved the final version of the manuscript and agree to be accountable for all aspects of the work.

## Supplemental Materials

**Supplemental Table S1**

Primer and RNA list used in the study.

**Supplemental Table S2**

Gene models located in the *b-4* linked genetic locus narrowed by ddRAD-seq analysis. FPKM values were obtained from RNA-seq analysis.

**Supplemental Table S3**

Final *b-4* candidate genes based on expression data. Mutations and False Discovery Rate (FDR) values at 24 and 48 hpo between p50T and e37 strain are listed.

**Supplemental Table S4**

Summary of results from CRISPR/Cas9-mediated knockout of the *KWMTBOMO12119*.

**Supplemental Fig. S1**

The absorption spectrum of ganglion pigments in (A) normal state, (B) oxidized state, and (C) reduced state. Ganglion and brain pigments were extracted from 20 larvae using 50 μL of 2% hydrochloric acid in methanol (v/v) and then diluted four times. After centrifugation (3000 × g for 5 min), supernatants were collected and used for analyses.

**Supplemental Fig. S2 Visualization of the *KWMTBOMO12119* gene transcripts using Sashimi plots.**

The gene structure of the *KWMTBOMO12119* is shown at the bottom. The arrow shows the direction of gene transcription. RNA-seq read counts obtained from eggs (0, 24, 48, and 72 hpo) are shown by red (e37 strain) and green (p50T strain) peaks on each exon. Numbers indicate RNA-seq reads spanning splice junctions.

**Supplemental Fig. S3 The insertion of a transposable element in *KWMTBOMO12119* gene in mutant strains e36 and No. 903.**

(A) Location of genomic PCR primers. Arrowheads indicate the position of forward and reverse primers, Bmmah_exon4to6_seq_F03 (F03) and Bmmah_exon4to6_seq_R02 (R02) listed in Table S1, respectively. The arrow shows the direction of gene transcription. (B) Gel electrophoresis of PCR products amplified from genomic DNA obtained from adult legs. (C) The genomic sequence around the insertion. A transposable element was inserted at the boundary between exon 5 and intron 5. Mutated nucleotides are shown in red letters.

**Supplemental Fig. S4 Pigmentation of larval brain and ganglions in *KWMTBOMO12119* knockout mutants.**

Representative larval brain and ganglions of *KWMTBOMO12119* knockout strain (*Δ4*). Scale bar: 1 mm.

**Supplemental Fig. S5 Egg pigmentation in the *os* knockout mutant.**

A representative *os* knockout egg. Scale bar: 1 mm. The *os* knockout strain (*os^In4^*) was derived from descendants of knockout individuals in Tomihara et al. (2019).

**Supplemental Fig. S6 Larval phenotype of the *Bmmah* knockout mutant.**

(A) A representative *Bmmah* knockout (*Δ4* strain; *Bmmah^Δ4^), os* knockout (*os^In4^*), and wild-type (p50T) larva. Scale bar: 1 mm. (B) Enlargement of the larval integument. The dorsal vein is visible beneath the surface of the integument in *os* knockout larva, but not in the wild-type and *Bmmah* knockout larva. Scale bar: 1mm.

